# Unravelling the genomic basis and evolution of the pea aphid male wing dimorphism

**DOI:** 10.1101/156133

**Authors:** Binshuang Li, Ryan D. Bickel, Benjamin J. Parker, Neetha Nanoth Vellichirammal, Mary Grantham, Jean-Christophe Simon, David L. Stern, Jennifer A. Brisson

## Abstract

Wing dimorphisms have long served as models for examining the ecological and evolutionary tradeoffs associated with alternative morphologies [1], yet the mechanistic basis of morph determination remains largely unknown. Here we investigate the genetic basis of the pea aphid (*Acyrthosiphon pisum*) wing dimorphism, wherein males exhibit one of two alternative morphologies that differ dramatically in a set of correlated traits that inclused the presence or absence of wings [2-4]. Unlike the environmentally-induced asexual female aphid wing polyphenism [5], the male wing polymorphism is genetically determined by a single uncharacterized locus on the X chromosome called *aphicarus* (“aphid” plus “Icarus”, *api*) [6, 7]. Using recombination and association mapping, we localized *api* to a 130kb region of the pea aphid genome. No nonsynonymous variation in coding sequences strongly associated with the winged and wingless phenotypes, indicating that *api* is likely a regulatory change. Gene expression level profiling revealed an aphid-specific gene from the region expressed at higher levels in winged male embryos, coinciding with the expected stage of *api* action. Comparison of the *api* region across biotypes (pea aphid populations specialized to different host plants that began diverging ~16,000 years ago [8, 9]) revealed that the two alleles were likely present prior to biotype diversification. Moreover, we find evidence for a recent selective sweep of a wingless allele since the biotypes diversified. In sum, this study provides insight into how adaptive, complex traits evolve within and across natural populations.

## Results and Discussion

### Mapping identifies the *api* locus

Winged and wingless male pea aphids (Figure 1A) are genetically determined by a single locus called *aphicarus* (*api*). This locus was previously localized to a 10cM region on the X chromosome (Figure 1B, top) [6], a chromosome estimated to contain a third of the genome [10]. To narrow down this region, we selfed F1 individuals from the *api* mapping line produced by this original study to create F2 individuals. Using a panel of 448 of those F2 pea aphids, we simultaneously identified and scored single nucleotide polymorphisms (SNPs) using multiplexed shotgun sequencing [11]. QTL analysis of these data resulted in the identification of 19 scaffolds containing SNPs with LOD scores higher than the 1% significance level of 7.6 generated by 1000 permutations (Table S1).

**Figure 1.**
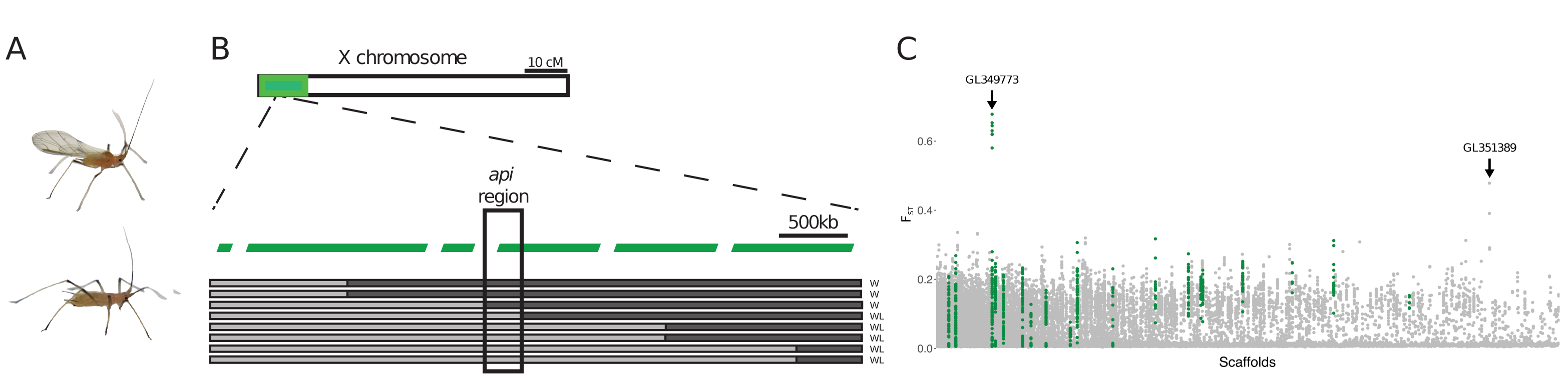
Linkage and association mapping of the *api* region. (A) Winged (top) and wingless (bottom) males. (B) The *api* region was initially localized to a 10cM region on the X chromosome (green box, top [6]). Seven ordered scaffolds in this region are shown in green. Further refinement of the linkage map narrowed the location of *api* to a ~130kb region spanning two genomic scaffolds (black box). The inferred recombination breakpoints of eight recombinant F2 individuals using RFLP markers are shown below the scaffolds. Black indicates sequence from the winged parent and grey from the wingless parent, and phenotype is indicated to the right (W:winged, WL:wingless). The recombination breakpoints are approximate (see methods for details). (C) Genome-wide view of genetic differentiation between winged and wingless males using F_st_ values calculated from 20kb windows across all scaffolds greater than 20kb (1,896 scaffolds). Scaffolds are ordered by their scaffold number, which is roughly from largest to smallest. F_st_ values from the 19 scaffolds identified from the recombination mapping analysis are indicated with green points.

Concurrently, to perform genome-wide association mapping, we sequenced the genomes of 44 pooled winged and 44 pooled wingless males collected from alfalfa (*Medicago sativa*) plants across the U.S. (Table S2) to a total of 68X and 70X coverage, respectively. F_ST_ analysis between the winged and wingless sequenced pools revealed two scaffolds with high levels of differentiation (Figure 1C). The first was a scaffold identified from the QTL analysis (LOD=17.5), while the second was not considered in the QTL analysis because it was smaller (42 kb) than the minimum scaffold size used in QTL analysis (scaffolds>100kb).

We physically ordered the 19 genomic scaffolds containing SNPs with LOD scores greater than 7.6, plus the smaller scaffold with a high F_ST_ value. For ordering, we assayed a restriction fragment length polymorphism (RFLP) marker for each scaffold using a panel of 40 F2 individuals that each carried a recombination event between the previous closest *api* flanking markers identified by Braendle et al. [6]. 16 of the 19 scaffolds were localized within these flanking markers, confirming the effectiveness of the QTL analysis. Two of these 16 scaffolds contained RFLPs with perfect association with all 40 F2s (see the *api* region noted in Figure 1B); these were the same two scaffolds identified from the F_ST_ analysis. Thus, recombination and association mapping each implicated the same two genomic scaffolds, a smaller scaffold (~42kb, GL351389) and a larger scaffold (>350kb, GL349773; this scaffold is misassembled after position ~350kb). We compared these scaffolds to their homologous regions from the peach-potato aphid (*Myzus persicae*) and Russian wheat aphid (*Diuraphis noxia*) genomes [12, 13] and determined that these two scaffolds sit proximately, in opposing orientation (Figure S1). Both of these species have a single genomic scaffold that spans the entire *api* region, with each species’ single scaffold containing homologous regions to the two pea aphid scaffolds. We developed additional RFLP markers in this region, which narrowed down the *api* region to between position 25kb on the smaller scaffold and position 107kb on the larger one, defining an approximately 130kb *api* region spanning the two scaffolds.

### High associations between SNPs in the *api* region and the winged and wingless males

The winged and wingless male pooled sequence (pool-seq) data highlighted many SNPs across the ~130kb *api* region that are strongly associated with the male wing phenotype (Figure 2A). The wide distribution of associated SNPs may indicate that this is a region of low recombination or that the *api* phenotype is generated by multiple SNPs that are maintained in linkage disequilibrium. The pool-seq data were generated from pea aphids collected from alfalfa. We wanted to determine whether a broader sample, across biotypes, would narrow down the causative polymorphism. As noted above, the pea aphid is actually a species complex with as many as 15 host plant adapted lineages, called biotypes, that began diverging 8,000-16,000 years ago [8, 9]. Biotypes have limited gene flow between them, and the ones with the least genetic exchange have been described as incipient species [8, 9]. We used the complete genomes of 23 genotypes from nine different biotypes (Table S3) to investigate patterns of association in the *api* region: nine winged allele carrying genotypes from five biotypes, and 14 wingless carrying genotypes from six biotypes (two biotypes, pea and alfalfa, had individuals of both allele classes). This data set thus overlapped with the alfalfa pool-seq data in that they both contained alfalfa biotype aphids, but allowed us to further narrow the informative genomic region. Because of the small sample size of this nine-biotype data set, here we focus exclusively on the SNPs that perfectly segregated with the winged and wingless males. There are 130 SNPs that meet that criteria. These 130 SNPs are indicated in orange, overlaid on the pool-seq association data (Figure 2A).

**Figure 2.**
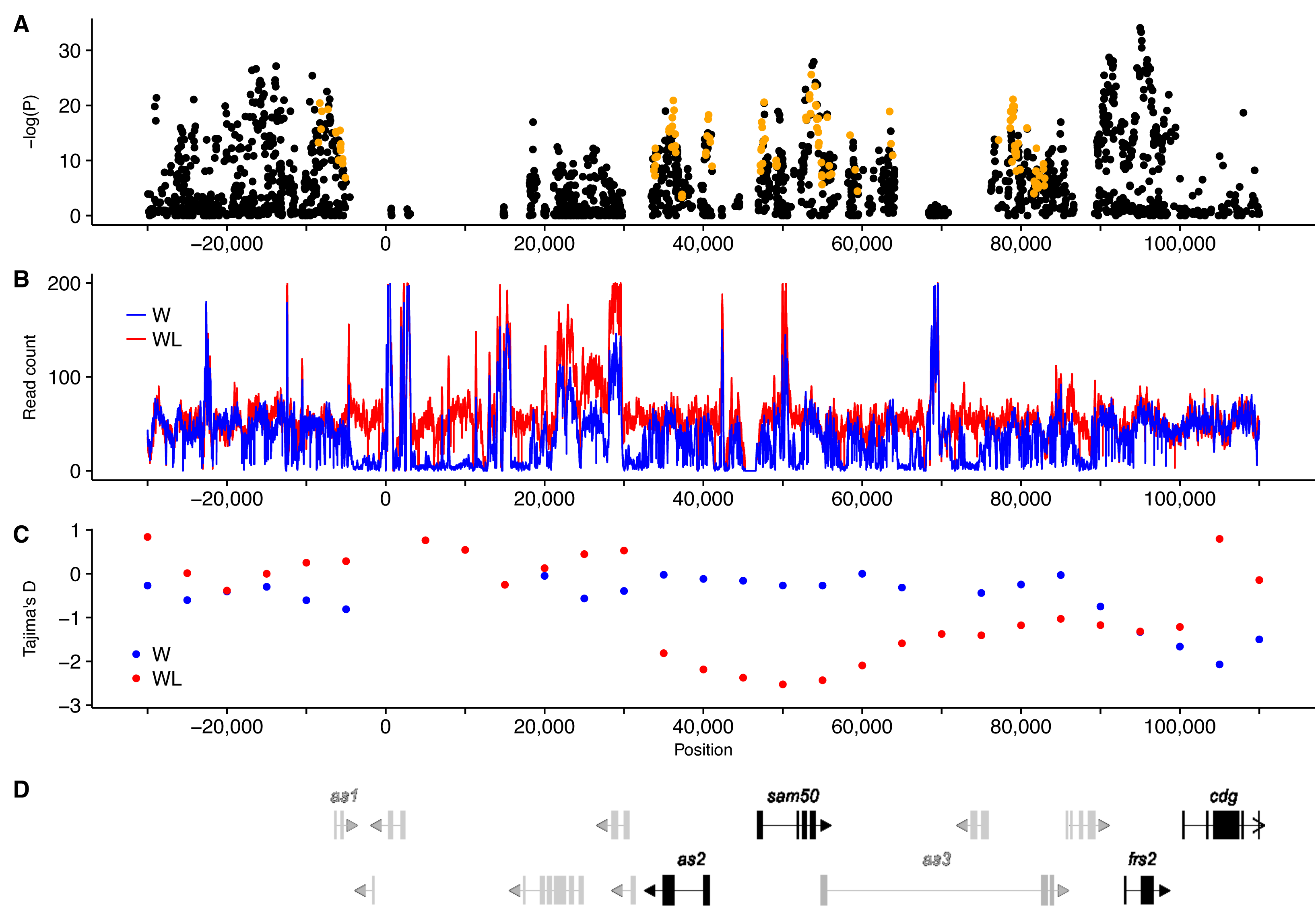
Population genetics statistics in the 130kb *api* region defined by recombination mapping. (A) Points show the large number of highly associated SNPs in the *api* region, illustrated as the -log(P-value) from a Fisher’s exact test between SNPs and the male wing phenotype using the alfalfa biotype pool-seq data. Orange points are SNPs that are perfectly segregating with the phenotype across the biotype data. 0 kb is the breakpoint between scaffolds GL351389, the smaller *api* scaffold is indicated by negative values, and GL349773, indicated with positive values. (B) The read count per site for pool-seq of winged and wingless males is shown on the y-axis, with a maximum of 200. The lower read depth in winged males despite near-equal sequence effort indicates sequence divergence. (C) The y-axis shows Tajima’s D values calculated across 10kb windows in 5kb steps for winged and wingless males separately. Only windows with greater than 30% of sites having a read count of at least 20 are presented. Some windows have insufficient data to calculate Tajima’s D. The low Tajima’s D values in the wingless males indicate a selective sweep. (D) Annotated genes in the *api* region. Grey indicates genes not expressed in any publicly available RNA-Seq data set, while black indicates expressed genes. *as1*, *as2*, *as3*=three aphid-specific genes, *sam50*=mitochondrial sorting and assembly gene, *frs2*=fibroblast growth factor receptor substrate, *cdg*=chromodomain-containing gene.

These two association studies, combined, indicate that the region that most likely contains *api* is the ~95kb region that includes the ~10kb end of GL351389 (the smaller *api* scaffold), and the first 85kb on GL349773 (the larger scaffold) (Figure 2A). Within this 95kb region, there are multiple regions with considerable sequence divergence such that the Illumina sequence reads from winged alleles cannot be aligned to the wingless reference genome, appearing as regions with low coverage from the winged pool (Figure 2B). These regions, from position ~5kb to ~33kb and from position ~65kb to ~75kb, could potentially be the causative variation, or contain the causative SNP(s). Across the whole genome, the winged and wingless pools were sequenced with near-equal effort (68X and 70x respectively), so sequence differences between the alleles are driving this pattern.

### A candidate gene at the *api* locus

The ~130kb *api* region contains 13 annotated genes in the pea aphid annotation, v.2.1 (Figure 2D; Table S4). The pea aphid genome annotation was primarily derived from gene prediction algorithms with aid from sequence information from female RNA libraries [14], leaving the possibility that male-specific genes in the region were missed during annotation efforts. We therefore sequenced four RNA-Seq libraries constructed from different stages of winged and wingless male embryonic cDNA, but did not detect any unannotated genes in the region. Of the 13 annotated genes, seven have transposable element-related annotations (Table S4). The remaining six genes code for three aphid-specific proteins with no conserved domains (*as1*, *as2*, and *as3*), a mitochondrial sorting and assembly gene (*sam50)*, a fibroblast growth factor receptor substrate (*frs2*), and a chromodomain-encoding gene (*cdg*) (Figure 2D). The seven TE-related genes, along with *as1*, have no discernable gene expression in our male embryo RNA-seq data or the 38 RNA-Seq libraries (36 from females of different ages, two from adult males) publicly available on <u>Aphidbase.com</u>. *as2*, *sam50*, *frs2*, and *cdg* all exhibited evidence of expression in our male-specific RNA-Seq libraries (Table S4); *as3* was not expressed in our male RNA-seq libraries, but was expressed in female libraries on Aphidbase [15]. We conclude these five genes (*as2*, *as3*, *sam50*, *frs2*, and *cdg*) are the only annotated genes that are transcribed and thus the only functional genes in the region.

We found no nonsynonymous changes that very strongly associated with the *api* phenotype across the pool-seq and biotype data; all nonsynonymous sites contained multiple reads in the pool-seq data that contradicted the association. We thus inferred that the mutation(s) that differentiates winged and wingless males must be regulatory. We measured the expression levels of the five genes (*as2*, *as3*, *sam50*, *frs2*, and *cdg*) that had evidence of transcription, using qRT-PCR. Winged and wingless males are morphologically different by the second nymphal instar [16] and wing morph determination in the environmentally induced wing polyphenism in pea aphid females occurs embryonically [17]. We thus reasoned that the action of *api* would occur embryonically, but that potentially the first nymphal instar may be important, too. Therefore, we focused on two developmental stages: embryos and first instar nymphs.

We observed that *as3* is not expressed in males at these stages, consistent with the RNA-Seq results. Among the four expressed genes (*as2*, *sam50*, *frs2*, and *cdg)*, only *as2* was differentially expressed between winged and wingless embryos (two-sided t-test, *P*=0.01; Figure 3), with two-fold higher expression in the winged embryos. No genes significantly differed in expression as first instars. *as2* is found in all sequenced aphid genomes [12, 13, 18], but is not found outside of aphids. *as2* is physically located in the center of our identified region near a large number of linked markers. It is also the only gene in the region that shows differential expression in the embryo stage, when *api* is likely to act. Thus, *as2* is the most likely candidate for *api*, although this hypothesis requires further validation.

**Figure 3.**
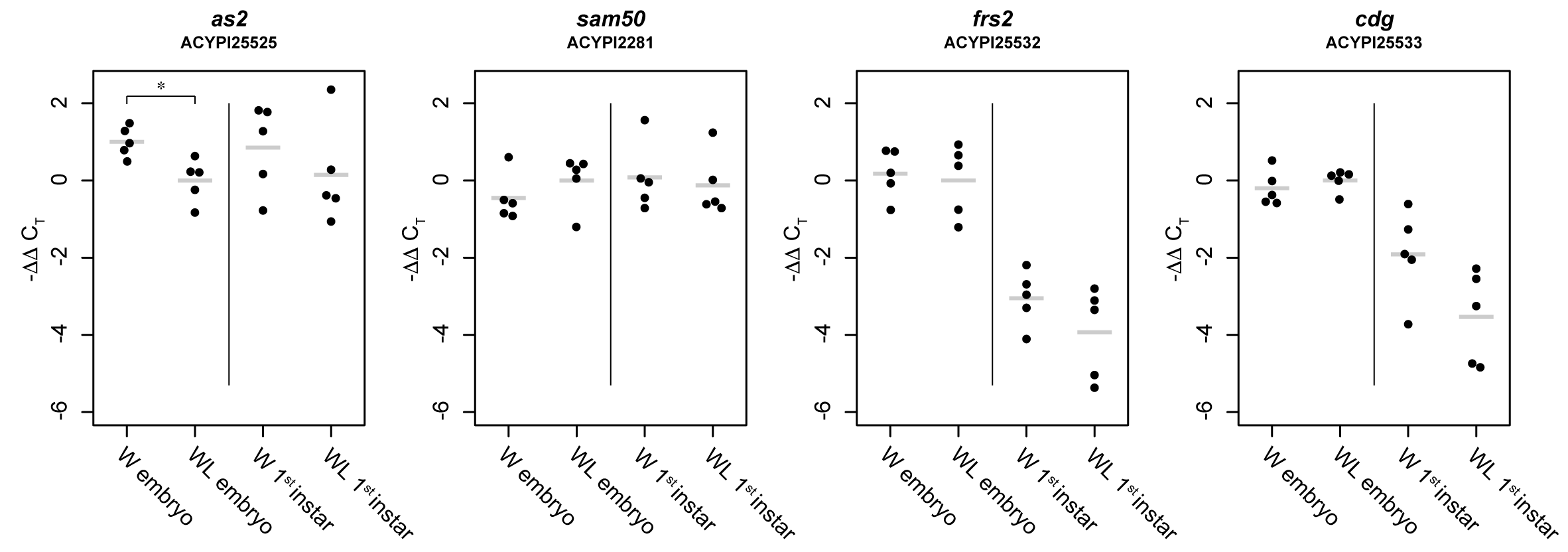
Gene expression levels for the four expressed genes in the *api* region. Each gene was measured in five biological replicates collected from winged and wingless male embryos and first instar nymphs. Each point represents a replicate. Y-axes show the ΔC_T_ values for each sample subtracted from the average ΔC_T_ value of wingless embryos (-ΔΔC_T_): higher values therefore represent stronger relative gene expression. Short grey horizontal lines show means. * indicates *P*<0.05.

### Molecular evolution at the *api* locus across pea aphid biotypes

To investigate the evolution of the *api* region within the pea aphid complex, we used the 23 resequenced genomes from nine biotypes. To determine the evolutionary history of the X chromosome in these lineages we constructed a phylogeny based on DNA polymorphisms from scaffolds located across the X chromosome, but not in the *api* region (Figure 4A). This analysis confirmed the genetic grouping of individuals from the same biotype, and that there is a continuum of sequence divergence across the complex of biotypes [9]. The winged and wingless phenotypes are scattered across this phylogenetic tree. In contrast, when we constructed trees from 10kb windows across the *api* region, we found complete separation of the winged and wingless genotypes (Figure 4B; Figures S2-3), suggesting that the wing morph is determined by the same variation in the different biotypes. Furthermore, the *api* region shows a different pattern of evolution than the rest of the X chromosome. These data suggest that the winged and wingless alleles pre-existed as standing variation before the biotypes split, and both alleles have been segregating in at least some lineages since then.

**Figure 4.**
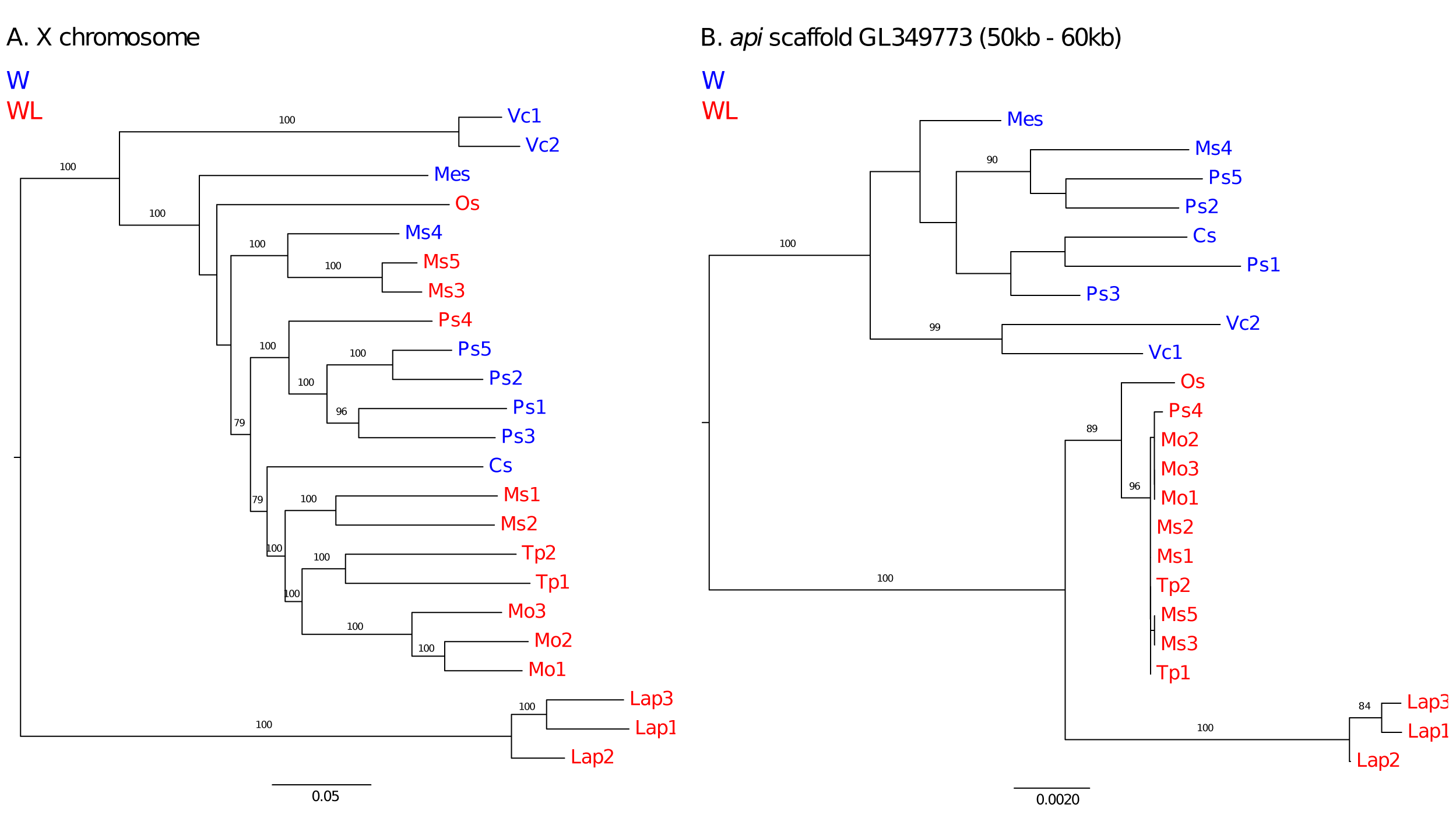
Relationships among winged and wingless genotypes from a range of pea aphid biotypes. (A) The evolutionary history of the X chromosome based on 27.6 Mb of sequence from across the X chromosome. (B) Phylogenetic tree based on positions 50-60kb in scaffold GL349773 in the center of the *api* region. Winged (W) lines are in blue, wingless (WL) in red. Trees are inferred using maximum likelihood, and numbers on the tree branches are bootstrap values; bootstrap values below 75 are not shown. Abbreviations: Mo: *Melilotus officinalis,* Tp*: Trifolium pratense*, Lap: *Lathyrus pratensis,* Ps: *Pisum sativum*, Ms: *Medicago sativa*, Os: *Ononis spinosa*, Mes: *Melilotus suaveolens*, Vc: *Vicia cracca*, Cs: *Cytisus scoparius.*

### A recent selective sweep of the wingless allele

The wingless alleles exhibit less genetic differentiation than the winged alleles in the *api* region (short branch lengths in Figure 4B; Figures S2-S3). To further explore this observation, we examined patterns of sequence variation in the alfalfa biotype using the pool-seq data. Since the two alleles do not seem to be freely recombining with one another, we examined Tajima’s D [19] separately in the two alleles to understand the evolutionary history of each. We found markedly lower Tajima’s D values in wingless males between positions ~35kb and ~65kb (Figure 2C), suggesting a selective sweep in the wingless allele. This signature of a selective sweep is located just upstream of *as2,* our *api* candidate gene. Within the *api* biotype trees (Figure 4B; Figures S2-S3), there is a long branch leading to three individuals of the *Lathyrus* (Lap1-3) biotype, a biotype that does not seem to hybridize with the other biotypes [9]. All examined biotypes currently carry both the winged and wingless *api* alleles ([20], Li et al., unpublished data). While the winged and wingless alleles arose before diversification of biotypes, it is not clear if both alleles have been maintained by natural selection in all biotypes since then. There is still limited gene flow between the biotypes [8] which could allow a biotype to recover an allele if it were lost. The reduced variation and the pattern of Tajima’s D in the wingless allele suggest that one wingless allele variant has recently swept through the alfalfa biotype (Figure 2C) and the other biotypes, except the reproductively isolated *Lathyrus* biotype (Figure 4). It is not clear if this is a new evolutionarily advantageous wingless allele (or a linked favorable allele), or the reintroduction of a wingless allele that has been lost from a large number of biotypes and recently become favorable. In addition, the selective advantage of this wingless allele is unclear but decreased dispersal due to the wingless phenotype could reinforce ecological speciation in the pea aphid complex by increasing mating within biotypes [21].

## Conclusions

Our study shows that the pea aphid provides a robust model for investigating the molecular basis of morphological variation. We have identified a single ~130kb-region of the pea aphid X chromosome that causes the differences between winged and wingless males. *as2*, an aphid-specific gene in the region, is expressed at higher levels in winged embryos relative to wingless embryos, making it a strong candidate for regulating this wing dimorphism. This study is the first demonstration of the genetic basis of a dispersal dimorphism, which have evolved repeatedly across insects because of the respective benefits of having winged and wingless morphs [1]. Moreover, we have demonstrated that a complex whole body phenotype can be regulated by what is likely a single gene. Finally, our results form the foundation for future comparative studies aimed at (1) discovering how male wing dimorphisms have been lost and gained repeatedly during the evolution of aphids [22, 23], (2) better understanding the factors promoting the maintenance of the male wing polymorphism, and (3) determining how the male wing polymorphism relates to the environmentally determined female wing polyphenism [24], which is present in most extant aphid taxa [3].

## Figure Legends

**Supplemental Figure 1. Comparison of scaffolds from three aphid species.** The pea aphid, *Acyrthosiphon pisum,* and *Myzus persicae* diverged from one another around 43 MYA [25]; *Diuraphis noxia* is more distantly related from both [26]. Lines are genomic scaffolds, with arrows indicating the orientation of a scaffold. Whole scaffolds from *D. noxia* and *M. persicae* are shown. The entire scaffolds of GL350308 and GL351389 and the homologous region of GL349773 from *A. pisum* are presented and to scale (GL349773 is misassembled above position ~350,000 and is not shown entirely).

**Supplemental Figures 2-3. Relationships among 9 winged and 14 wingless genomes from across nine biotypes.** Unrooted trees are constructed from nucleotide variation from 10kb windows in the *api* region. Information below each tree indicates the scaffold, the 10kb window on the scaffold used, and the number of variable sites in that window. Trees for intervals containing less than 150 variable sites are not shown. Individuals containing *api* winged alleles are colored in blue and wingless alleles in red. Numbers on the tree branches are bootstrap values. Abbreviations: Mo: *Melilotus officinalis,* Tp*: Trifolium pratense*, Lap: *Lathyrus pratensis,* Ps: *Pisum sativum*, Ms: *Medicago sativa*, Os: *Ononis spinosa*, Mes: *Melilotus suaveolens*, Vc: *Vicia cracca*, Cs: *Cytisus scoparius;*

## Materials and Methods

### Linkage mapping

Braendle et al. [6] previously established an *api* linkage mapping population. We used the F1 line from that population to generate additional F2 recombinants. Specifically, F1 asexual females were placed on *Vicia faba* plants in an incubator at 16°C and a photoperiod of 12h light and 12h dark. After two generations of asexual reproduction, these conditions induce the production of sexual females and males. We crossed F1 females to F1 males, collected fertilized eggs, sterilized them in 1% calcium propionate on Whatman paper, and placed them in an incubator that alternated between 4°C for 12h light and 0°C for 12hr dark. After 90 days, eggs were removed from this incubator and placed in a 19°C incubator that alternated between 16h light and 8h dark. F2 hatchlings were transferred to individual plants for asexual reproduction to establish a line of that F2 individual.

Genomic DNA of 448 F2 females and 22 F2 males were isolated, quantified, and diluted to 2ngs/μl. Multiplexed shotgun sequencing of all 470 F2 individuals was carried out according to [27] with minor modifications: individual F2 DNA quantity for Mse1 digestion was increased to 10ngs and the unique barcoded adapter concentration was reduced to 2.5nmols. These changes were made to increase ligation efficiency and to reduce the formation of ligated linker-dimers. The final amplified libraries were sequenced on an Illumina Genome Analyzer.

We identified genomic scaffolds that exhibited linkage with *api* using Rqtl [28]. For each scaffold of interest, we developed a diagnostic restriction fragment length polymorphism (RFLP) marker. RFLP markers were tested on a panel of 10 F2 individuals to confirm that the scaffold was indeed X-linked and linked to *api*. The scaffolds were ordered relative to one another using a panel of up to 40 F2 individuals. Recombination breakpoints for the two scaffolds in the *api* region were localized to within ~10kb (left side defined by markers at positions 15,767 and 25,460 on GL351389 and right side by markers at positions 97,258 and 107,073 on GL349773). The others scaffolds contained only one RFLP marker each and thus breakpoints in Figure 1C are approximated by representing them in the middle of the adjacent scaffolds.

### Pool-seq

We used 44 winged and 44 wingless male pea aphids induced from females collected from Nebraska, New York, California and Massachusetts (Table S2). Genomic DNA (gDNA) from each male was extracted using the Qiagen DNeasy Blood & Tissue Kit. 50ng of gDNA from each male was pooled together with males of the same phenotype. Paired-end libraries were prepared with the TruSeq DNA PCR-Free Library Preparation Kit at the University of Rochester Genomics Research Center and sequenced on an Illumina HiSeq2500 Sequencer with paired 125nt reads. Reads were mapped to the pea aphid reference genome v.2 using bwa (v.0.7.9a-r786) [29] using default parameters. Reads were filtered for a mapping quality of 20 and BAM files were sorted by coordinates using samtools (Version: 0.1.19-44428cd). The coverage of both libraries was calculated using the samtools depth function for the mapped reads.

### Association, F_ST_, and Tajima’s D analyses

An mpileup file was created from the winged and wingless male BAM files using the samtools mpileup function with the -B option to disable BAQ computation. This was further processed by the mpileup2sync.jar script in PoPoolation2 v.1.201 [30] to generate a synchronized mpileup file (sync file), with fastq type set to sanger and minimum quality set to 20. For association analyses, Fisher’s exact tests were performed using the <u>fisher-test.pl</u> script included in PoPoolation2 with a window-size of 1, step-size of 1, minimum coverage of 2, and minimum allele count of 2. F_ST_ values were calculated with the <u>fst-sliding.pl</u> script included in PoPoolation2 with a window size set to 20kb, a step size set to 10kb, minimum coverage set to 30, a maximum coverage set to 200, pool size set to 44. The <u>variance-sliding.pl</u> script was used to calculate Tajima’s D, setting a minimum allele count of 2, a minimum base quality of 20, a minimum coverage of 10, a maximum coverage of 400, a pool size of 44, a window size of 10,000, and a step size of 5,000; fastq type was set to sanger. Data are shown for windows with a 30% minimum covered sequence fraction.

### Fully re-sequenced genomes

In addition to the reference pea aphid genome which is *api* wingless homozygous, we obtained the sequence of 22 additional pea aphid genomes: 9 carrying only the winged allele and 13 the wingless allele (Table S3). These genotypes have been collected in the wild as parthenogenetic females, on distinct legume species. Biotypes of these genotypes have been assigned based on their microsatellite profiles [9] and representatives have been then selected for genome resequencing [31]. Lines were sequenced to 17X to 30X coverage with Illumina 100nt paired-end reads. The reads of each sample were aligned to the pea aphid reference genome with default settings in bowtie2 (version 2.2.1) [32]. The consensus sequence of each sample was acquired using the recommended pipeline in samtools (version 1.3.1) and bcftools (version 1.3).

### Phylogenetic trees

Trees were built using Raxml v. 8.2.9 [33] using the substitution model GTRGAMMA. One thousand bootstraps were run on distinct starting trees. For the neutral tree, we used a concatenated data set of genomic scaffolds from the pea aphid genome build v.2 with a probability of being on the X above 90% (111 scaffolds, total of 27.6 Mb [34]). The *api* trees were constructed using the 9 winged and 14 wingless biotype sequences in 10kb intervals. The trees were graphed using Figtree (v 1.4.0)[] and rooted by the included midpoint method.

### Male embryo RNA-seq

Stage 18 embryos [35] and a mixed sample of embryos younger than stage 18 were obtained separately for F2 lines homozygous for the *api* winged or wingless allele (four libraries total). Total RNA was isolated using TRIzol^®^ (Thermo Fisher Scientific, Waltham, MA). Library construction was performed using the TruSeq Stranded mRNA Sample Preparation Kit (Illumina, San Diego, CA). The four libraries were constructed and sequenced with single end 100 nt reads in one lane of an Illumina HiSeq 2500 sequencer at the University of Rochester Genomics Research Center. The reads were aligned to the reference genome using bowtie2 [32] and processed into bam files using samtools with default settings. Bam files were visualized with IGV (v.2.3.72) [36]. The coverage of mapped reads was reported using samtools and compared to the gene annotations. No coverage of more than six reads was discovered outside of annotated exon regions within the api region.

### Quantitative reverse-transcriptase PCR (qRT-PCR)

We also used the two *api* homozygous F2 lines to collect male embryos and first instar nymphs. To produce males, we transferred asexual female adults into an incubator at 16°C with a photoperiod of 12h light and 12h dark. Two generations in this environment resulted in asexual females whose offspring would be males and sexual females. We dissected stage 18 [35] embryos from the females or collected first instar nymphs. We confirmed the sex of embryo or nymph using an RFLP on the X chromosomes (forward primer: ATCGATGCTTTTGAATTGTTTTAC; reverse primer: TGTAGGGTCTCTTGAAGTTGTTTG; restriction enzyme: Taq*α*I; double bands are females while single bands are males) while simultaneously collecting tissue for RNA as in [37]. Five biological replicates were included. Each embryo replicate contained 20 embryos from 6 to 10 females, while each first instar nymph replicate contained 10 individuals produced by 3 to 5 females. Quantitative PCR was performed on a Bio-Rad CFX-96 Real-Time System using 12μL reactions of 40ng cDNA, 1X PCR buffer, 2nM Mg+2, 0.2nM dNTPs, 1X EvaGreen, and 0.025 units/μL Invitrogen Taq with the following conditions: 95°C 3 min, 40x (95°C 10s, 55°C 30s). Primer concentrations were optimized to 100+/-5% reaction efficiency with an R^2^ value of > 0.99 [G3PDH (ACYPI009769): 400nM Forward primer, 350nM Reverse primer; NADH (ACYPI009382): 350nM F, 300nM R primer; 2281: 175nM; 25525: 150nM; 25532: 250nM; and 25533: 150nM]. Each of the five biological replicates was run on a single plate, with three technical replicates of each reaction. ΔCt values were calculated by subtracting the average C_T_ value of the two endogenous controls (G3PDH and NADH) from the C_T_ values of each target gene. For each pair of winged and wingless samples, ΔC_T_ values were analyzed using two-sided t-tests after checking for normality.

## Acknowledgements

We gratefully thank Jen Keister for technical assistance. This research was supported by award R00 and R01GM116867 from the National Institute of General Medical Sciences to J.A.B. RNA-Seq and pool-seq sequencing was performed at the University of Rochester Genomics Research Center. J.C.S. was supported by the Agence Nationale de la Recherche (grant ANR-11-BSV7-007). Sequencing of the YR2 line was performed at the Genoscope (project 62 AAP 2009/2010 to J.C.S.). We thank Prof. Shin-Ichi Akimoto who provided sample and *api* phenotype for pea aphid lines 09003A, Sap05VC7 and Iwamizawa.

## Author Contributions

Conceptualization, J.A.B., R.D.B., and D.L.S.; Methodology, J.A.B., R.D.B., and D.L.S.; Formal Analysis: B.L., R.D.B., and D.L.S.; Investigation, B.L., R.D.B., B.J.P., N.N.V., M.G., D.L.S., and J.A.B.; Writing – Original Draft, J.A.B., R.D.B., and B. L.; Writing – Review & Editing, D.L.S. and J.-C.S.; Funding Acquisition, J.A.B.; Resources, J.-C.; Supervision, J.A.B.

